# Precursors of Sea Star Wasting: Immune and Microbial Disruption During Initial Disease Outbreak in Southeast Alaska

**DOI:** 10.1101/2025.07.05.662347

**Authors:** Andrew R. McCracken, Aly Rodger, Chanchal Saratkar, Sage Sularz, Sophia Brusch, Joaquin C. B. Nunez, Melissa H. Pespeni

## Abstract

Sea Star Wasting Disease (SSW) has devastated sea star populations along the North American Pacific coast since 2013, yet the mechanisms of disease progression, particularly in natural environments, remain unclear. Here we integrate transcriptomic and microbial data from wild *Pycnopodia helianthoides* sampled across sites affected and unaffected by SSW in southeast Alaska during the initial outbreak recorded in the region in 2016. Individuals exposed to SSW but lacking visible symptoms showed elevated expression of complement system components, pathogen recognition genes, immune regulatory and cell death pathways. Alongside signs of immune activation, genes involved in maintaining extracellular matrix composition, tissue remodeling, and cell adhesion were differentially expressed, indicating early disruption of tissue homeostasis preceding visible wasting symptoms. Gene ontology analysis revealed enrichment of immune response, cell-cell adhesion, response to oxygen levels and nervous system regulatory pathways. Furthermore, network analyses revealed differentially abundant microbes in Exposed individuals—notably *Vibrio spp*.—were highly correlated with immune response, tissue integrity, stress and detoxification genes in network modules. Together, our findings offer insight into early host-pathogen dynamics in wild populations, underscoring putative links between immune activation and microbial community shifts with the onset of SSW disease.

## Introduction

Infectious diseases, both endemic and emerging, have increased in frequency and severity over the past few decades and are projected to further increase over time (1–5). While the general consensus is that anthropogenic drivers amplify these patterns, identifying the direct factors influencing the emergence of wildlife epidemics is often difficult (6, 7). This is especially true in marine ecosystems where changes in temperature, ocean chemistry, pollution, and habitat destruction can compromise wildlife health in unpredictable ways (8–10). Rapid change to environmental norms may not only cause stressful conditions for marine organisms, but also favor the spread of disease (3, 8, 11). Due to these complex biotic and abiotic interactions, pinpointing the causes of disease necessitates an integrative approach that combines genomics, microbial ecology, and environmental context to understand the driving factors of emerging diseases.

Sea Star Wasting (SSW) disease is one of the largest emerging marine epizootics ever recorded due to its large geographic range and high mortality rates (12–14). Yet, despite its unprecedented scale, the cause(s) of SSW remain under investigation, with growing evidence pointing to microbial agents as a potential driver of disease (15). Since the summer of 2013, a series of recurring outbreaks have devastated sea star populations along the west coast of North America impacting over 20 species with mortality rates as high as 100% for some species at some sites (12–14, 16). SSW has expanded its range north in the Pacific each year with evidence that it is reaching the Atlantic in both North American and European waters (17, 18). Many sea star species are integral members of their ecosystems and their removal has caused the collapse of ecological communities, resulting in the loss of biodiversity with cascading ecological and economic consequences (13, 14, 19–21).

A significant obstacle impeding this field is identifying the causative agent(s) of disease. SSW advances quickly, with mortality progressing as soon as 24 hours after the first visible signs of disease (14). This rapid progression from visibly healthy to death makes it difficult to study in the lab and even more difficult in wild populations as it provides limited and time-sensitive opportunities for sample collection and monitoring. Efforts to connect the outbreaks to common disease driving factors such as temperature, density dependence, and or pathogenic agents have proven challenging (12, 22–25). Recent advances are beginning to clarify the microbial mechanisms involved. Consistent shifts in microbial community’s structures preceding and following signs of disease progression across multiple species (26–30). These changes are characterized by an increase in facultative and obligatory anaerobic microbes at the animal-water interface that thrive in low-oxygen environments rich in organic matter — such as a wasting sea star. It is hypothesized that the proliferation of anaerobic microbes on the surface of the sea star may reduce oxygen availability for the host, reducing respiration and increasing stress that may lead to a cascade of unknown factors driving the disease (28). A commonality across these studies has been the observed increased in abundance of *Vibrio* species associated with disease states and progression, a genus of bacteria commonly associated with other echinoderm related diseases such as tissue necrosis in the green sea urchin (31), Skin Ulceration Disease (SKUD) in the sea cucumber *Holothuria scabra* (32), and induced wasting-like disease in crown of thorns sea stars (33, 34). However, how these microbial shifts impact host physiology—and whether changes to the microbiome contributes to disease progression or is instead a consequence of disrupted homeostasis—remains unclear.

The sunflower sea star (*Pycnopodia helianthoides*) offers a powerful system for understanding host-microbiome interactions during marine disease outbreaks. As one of the largest and most ecologically impactful sea stars in the North Pacific, its dramatic decline due to SSW has provided a rare opportunity to investigate both host responses and microbial changes induced by natural disease exposure. Despite growing recognition of the importance of host-microbiome interactions in development, immunity, and behavior (35–37), most SSW studies have examined host transcriptomics or microbial shifts independently, rarely integrating both perspectives. Moreover, many have compared individuals that may have already been exposed to the disease, as samples were often collected from regions with ongoing or recent outbreaks (26, 28, 30, 38).

For example, in *Pycnopodia*, transcriptomic studies have revealed activation of immune, neural, and tissue remodeling genes associated with wasting symptoms (26, 38), providing foundational insight into the host response. However, these studies did not assess microbial dynamics concurrently, leaving the role of the microbiome in shaping or responding to disease unresolved. More integrative approaches in *Pisaster ochraceus* have begun to address this gap: recent work by Pespeni and Lloyd (2023) tracked host gene expression alongside microbial community shifts across disease stages, revealing associations between differentially abundant microbes and host genes involved in cell adhesion, defense, and collagen proteolysis. They also observed upregulation of genes related to low-oxygen conditions, such as hypoxia-inducible factor 1-α, supporting hypotheses that microbial activity may exacerbate disease through localized oxygen depletion (28–30). Yet it remains untested whether such host-microbe responses are conserved across species and truly reflect natural patterns of exposure in the wild. Ideally, we would compare gene expression and microbial profiles from animals in regions free from disease to exposed individuals (but not yet wasting), sampling the same tissues and sequencing both host-response and microbe abundance in tandem to understand the initial response to disease and microbiome disruption.

In this study, we analyze a unique set of samples collected from wild *P. helianthoides* in southeast Alaska taken in the summer of 2016 when the disease was first detected in the region. In a previous study, we identified signs of microbial dysbiosis preceding signs of SSW disease and progressing through disease onset in these wild populations of *P. helianthoides* (29). To better understand the relationship between host and microbiome in the progression of the disease in the wild, we analyze a subset of these samples measuring host transcriptomic and microbial abundances in naïve (i.e., healthy and no historical record of disease) and exposed (i.e., believed to be exposed the disease for the first time by cohabitating with wasting individuals) populations. Here, we generate host transcriptome data and integrate with previously generated 16S rRNA sequence data from the same individuals and tissue samples from wild *P. helianthoides* populations. We compare epidermal biopsies of sea stars sampled from sites with no evidence of wasting disease (i.e., “Naïve”; n =5) and from apparently healthy individuals in areas of active wasting disease (i.e., “Exposed”; n = 5)**. Based on previous research and preliminary observations, we developed and tested the following *a priori* hypotheses:** (**1**) that host gene expression would show activation of immune-related pathways, including inflammatory and antimicrobial resistance genes, prior to visible disease signs, and (**2**) that increased abundance of putative pathogenic microbes would correlate with the activation of host immune-related pathways, implicating certain microbes in the progression of SSW disease.

## Methods

### Sample Collection

Epidermal tissue biopsies of *P. helianthoides* were collected during the summer of 2016 off the coast of Southeast Alaska. As part of a previous study investigating the microbial communities associated with SSW in the wild, samples were collected using a ship and SCUBA diving at seven sites: two naïve sites with no recent or historical records of SSW, and five Impacted sites exhibiting their first recorded outbreak (29). From the naïve sites, 47 individuals were sampled. From the Impacted sites, 67 individuals were sampled — 20 appearing outwardly healthy despite being collected from outbreak locations (designated as “Exposed”), and 18 exhibiting visible signs of SSW (designated as “Wasting”). In total, 85 individuals were sampled across all sites. In the present study, we randomly selected five individuals using a random number generator from each group for transcriptomic and metagenomic analyses. After RNA sequencing, it was determined that the transcripts acquired from the wasting samples were not of high enough quality and were dropped from the study. Similarly, after DNA sequencing for metagenomics, it was determined that most of the sequence data was from *P. helianthoides*, thus the metagenomic aspect of the study was dropped. RNA sequencing data from the remaining 5 naïve and 5 exposed (N=10) individuals passed quality tests and were used for transcriptomic analyses.

### RNA Extraction, library preparation, and sequencing

RNA was extracted from each biopsy using a modified TRIzol protocol (TRIzol reagent ThermoFisher Scientific, Waltham, MA). Tissue was first lysed in 250 µl of TRIzol, followed by homogenization for 20 minutes using a plastic pestle and an additional 750 µl of TRIzol, with agitation on a Vortex Genie2 (Scientific Industries, Bohemia, NY). RNA extraction was performed by adding 200 ul chloroform, inverted 15 times, incubated for 3 minutes at room temperature (RT), and centrifuged at 4°C for 15 minutes at 12,000×g. The RNA-containing supernatant was transferred to a new tube and the previous step was repeated. 500 ul isopropanol and 1 ul 5 mg/ml glycogen was added then incubated for 10 minutes at RT. Samples were then centrifuged for 5 minutes at 7500×g at 4°C to precipitate the RNA from the supernatant. Samples were left to dry for 10 minutes at RT, then the RNA pellet was dissolved in 50 ul nuclease-free water. RNA quality and quantity was measured with a NanoDrop 2000 Spectrophotometer (ThermoFisher Scientific, Waltham, MA) and Qubit 3.0 Fluorometer (Life Technologies, Carlsbad, CA).

***Host RNA*** library preparation and sequencing was performed at Novogene (Novogene, Sacramento, CA facility). Libraries were prepared using poly-A selection and sequenced on an Illumina NovaSeq using 2 x 150 bp overlapping paired end reads, generating ∼40 million read pairs (∼6GB raw data) per sample. In the present study, ***Genomic DNA*** was originally collected for metagenomic purposes, however, sea star DNA swamped microbial DNA, thus we were able to use the sea star data to test for signatures of population structure between the two sampled sites using principal component analysis (PCA) and Admixture (see Supplemental methods). Genomic DNA was extracted from animal tissue samples using the DNeasy Blood & Tissue Kit (Qiagen, Hilden, Germany) according to the manufacturer’s protocol. Briefly, approximately 25 mg of tissue was lysed with proteinase K and Buffer ATL at 56 °C overnight, followed by DNA binding in the presence of Buffer AL and ethanol. The lysate was loaded onto spin columns, washed with Buffers AW1 and AW2, and eluted in Buffer AE. DNA concentration and purity were assessed using a Nanodrop spectrophotometer (Thermo Fisher Scientific). Genomic DNA library preparation and sequencing was also performed by Novogene on an Illumina NovaSeq platform using 2 x 150 bp overlapping paired end reads, generating ∼40 million read pairs (∼6GB raw data) per sample. Due to the poor representation of microbial DNA from host tissue, ***Microbial 16S rRNA*** data was collected to analyze patterns of taxonomic change among samples. Although metagenomics analysis is preferred for accurate species or strain-level microbial analysis, 16s sequencing may be more accurate and affordable to examine broader patterns of microbial diversity (39). Microbial 16S rRNa were generated by amplifying the V3 and V4 region of the 16S bacterial gene using the same extractions from the host RNA prep. 16S library prep and sequencing was performed at the Cornell Biotechnology Resource Center (Ithaca, NY) using 2 × 300 base pair overlapping paired-end reads on an Illumina MiSeq platform. For full details on 16S amplification and library prep, see McCracken et al 2023.

### Host RNA Mapping and Differential Abundance and Annotations

#### Read cleaning and mapping

Raw sequences yielded between 20-27 million paired end reads (average: 23.9M reads) per sample. Raw reads were trimmed and filtered for quality using fastP v0.23.4 (40), applying a Phred quality threshold of >30 and removing adapter contamination. After quality filtering, between 20-26 million reads (average: 23.0M reads) per sample remained. Quality metrics (e.g., base quality, GC content, and sequence length distribution) were visualized using the fastP summary report. Cleaned reads were aligned to the 24,184 nucleotide sequences for all the predicted genes of the *P. helianthoides* reference genome (augustus.hints.codingseq; (41)) and mapped read counts were obtained using `STAR` v2.7.11b (42) with default parameters to obtain a sample-to-gene-counts matrix for differential expression analysis. Samples from Naïve and Exposed individuals had an average of 82.5% (range: 75-87%) uniquely mapped reads to the reference. In contrast, samples from wasting individuals had poor and inconsistent mapping rates with an average of 26.8% (range 6-83%) uniquely mapped reads and were dropped from analysis.

#### Differential abundance

DESeq2 v1.40 (43) was used to normalize counts to account for variation in library size across samples and test for differential expression among host genes in `R` v3.4.2. Low-abundance genes were filtered out by removing transcripts which had fewer than 10 counts per sample on average. Out of the ∼24,184 mapped genes, 15,220 (62.9%) remained after keeping those with greater than an average of 10 counts per sample. Models were run in DESeq2 to test for differential expression between Naïve and Exposed groups as [MODEL: ∼ 0 + health_site_status]. An intercept-free model was used to estimate the difference of mean expression for each group, then compared against an intercept-inclusive model yielding the same results. The model aimed to capture early changes in the host’s physiological state upon disease exposure. All genes were annotated using Diamond’s Blast-X aligner (44) using the non-redundant NCBI database (45). A fasta file containing all genes was blasted for sequence similarity and the top annotations were returned for each gene. Gene names, blast annotations, and log2-fold change for all genes available on Dryad (**DOI: 10.5061/dryad.t1g1jwtfh**). In addition to recording the most differentially expressed genes between disease states, subsets of genes related to tissue homeostasis, nervous system function, cellular stress, and immune response, based on gene ontology mapping (described below) to identify early changes to these systems in response to disease exposure.

#### Gene Ontology

GO categories for all genes were obtained by mapping PANTHER (Protein Analysis Through Evolutionary Relationships) classifications available from the reference genome (41) to corresponding GO annotations. PANTHER classifications are maintained up to date with GO annotations as part of the PANTHER 19.0 database (46, 47). 18,324 genes were identified with PANTHER annotations from the reference genome and were mapped to corresponding GO categories using the PANTHER HMM Classification system available in the PANTHER database. To test if gene categories of specific biological functions were enriched in our differentially expressed genes, we performed gene ontology (GO) enrichment analysis with TopGO using the parent-child algorithm with Fisher-Exact method for enrichment scores (48). This was done for two groups of genes, those that were significantly upregulated in Exposed stars (down in naïve samples) and those that were downregulated in Exposed stars (up in naïve samples) where members within the group were assigned a binary value 1 or 0 representing differential expression or not, respectively. GO enrichment on Weighted Gene Correlation Network Analysis (WGCNA) modules (described below) were calculated in the same manner with a 1 representing within-module membership and 0 for no presence within the selected module. Categories with fewer than 5 genes or over 500 were dropped to avoid categories that were too small or too broad. For visualization, parent terms were gathered from each term found to be significantly enriched, and if that parent term was also significant, the child was dropped to keep the broader of the two categories to understand and visualize the higher-level processes at work. For the full list of enriched GO terms – including all parent and child terms – reference Tables S1 and S2.

GO terms were used to count and visualize the number of differentially expressed genes with biological functions related to immune response, nervous system function, stress response, and tissue integrity. These categories were chosen based on previous studies reporting symptomology and gene expression of actively wasting sea stars and our goal was to visualize changes to these genes in the early stages of disease exposure (12–14, 30, 38). We used a custom set of search terms to build a list of genes that had an associated GO function containing any of the following key terms for: immune functions (i.e., “immune”, “defense”, “antigen”), nervous system function (i.e., “nerve”, “nervous”, “neural”, “neuro”, “synapse”, “synaptic”, “glia”, “dendrite”, “dendritic”, “axon”), stress (i.e., “stress”, “heat shock”, “hsp”, “DNA damage”, “DNA repair”, “apoptosis”), and tissue integrity (“collagen”, “extracellular matrix”, “ECM”, “metallopeptidase”, “cell adhesion”, “cell-matrix adhesion”, “wound healing”, “tissue homeostasis”). Some more specific terms may be missed with this search approach, however, genes with a more specific GO annotation would also contain the broader category, and thus we believe we captured most genes in these categories.

### Microbial Differential Abundance and Annotations

Our previously generated 16S data (29) were reanalyzed using the 10 focal samples of the present study with the latest programs, packages, and databases. Qiime2 v.2024.5 (49) was used to obtain microbial abundance counts and classifications. Paired-end 16S reads were imported into Qiime2 and read quality assessed and visualized. Read quality was determined by Phred score (>30) then subsequently denoised and trimmed using DADA2 (50). DADA2 parameters were set at trimming forward reads at 16 bp and truncating at 289 bp and reverse reads were trimmed at 0 bp and truncated at 257 bp. Cleaned ASV’s were then classified using Qiime2 Naïve-bayes classification plugin against the Silva-138.1 non-redundant 99 SSU database (51). The Sylva database was formatted for Qiime-2 using RESCRIPt (52). Sylva reference v.138.1 sequence and taxonomy files were downloaded, reverse transcribed to cDNA format, low quality sequences removed, dereplicated, V3-V4 amplicon regions extracted, and used to train Qiime-2’s Naïve-bayes classifier. A counts matrix was exported containing ASV’s classified into observable taxonomic units (OTUs) at species-level, or as close to species as the classification allowed (such as genus or family). AncomBC-2 (53, 54) was used to calculate differentially abundant taxa between Naïve and Exposed (FDR *P*_adj_ < 0.05). We identified 252 unique observable taxonomic units (OTUs) present in our samples (**DOI: 10.5061/dryad.t1g1jwtfh**).

### Host-Gene and Microbial-Abundance Network Analysis

To identify potential host-microbe interactions, we used the WGCNA v.1.7.3 package in R to identify covarying host gene expression and microbial abundance in modules, i.e., networks (55). We first removed low abundance transcripts and OTUs that were missing from >70% of the samples in each respective dataset. DESeq2 was then used to normalize microbial abundance and gene expression count datasets separately to account for differences in read-depth across samples. Both datasets were then concatenated and log_2_ normalized to bring both datasets to a comparable scale suitable for joint analysis (***Figure S1***). Modules were constructed to identify genes and microbes that are “expressed” or appear in similar abundances in the combined dataset blind to site-health identity. To identify correlations between modules and site-health status, we ran Pearson’s correlation tests between Naïve and Exposed groups and their WGCNA module eigen-values. Modules most strongly correlated with Naïve or Exposed groups (*P* < 0.05; two-tailed *t*-test) were selected and examined for gene and microbe module membership. Genes belonging to highly correlated modules that contained both genes and microbes were then analyzed for gene ontology enrichment to discern functional categories associated with the modules. To identify highly correlated genes and microbes interacting within modules, we pruned module networks around focal genes or microbes of interest (such as *Vibrio*). Within each module network, genes and microbes are the nodes, while the correlation value between each is an “edge” that connects two nodes. Because all genes and microbes are connected in each module, we “pruned” the network to keep those with correlations greater than |0.5| and focus on connections between nodes of interest.

## Results

### Global transcriptomic and microbial profiles separate naïve and exposed groups in multidimensional space

Our transcriptomic analysis revealed that 2,006 genes (12.7% of the 15,743 tested) were differentially expressed between naïve and exposed sea stars (*P*_adj_ < 0.05 via Benjamini-Hochberg) with 743 genes (4.8%) upregulated in naïve individuals and 1,263 genes (8.0%) upregulated in Exposed individuals. Using global gene expression profiles, PCA revealed clear clustering of individuals by exposure status along PC 2 with 19% of the variance explained (**Figure 1A**). Our microbial analysis uncovered 39 (15.5%) OTUs that were differentially abundant between Naïve and Exposed groups, 10 of which decreased and 29 increased in abundance in the Exposed group relative to Naïve (FDR *P_adj_* < 0.05). PCA on microbial abundance profiles also distinctly separated naïve and exposed groups with 38% of the variance explained by PC 1 (**Figure 1B**).

**Figure 1.**
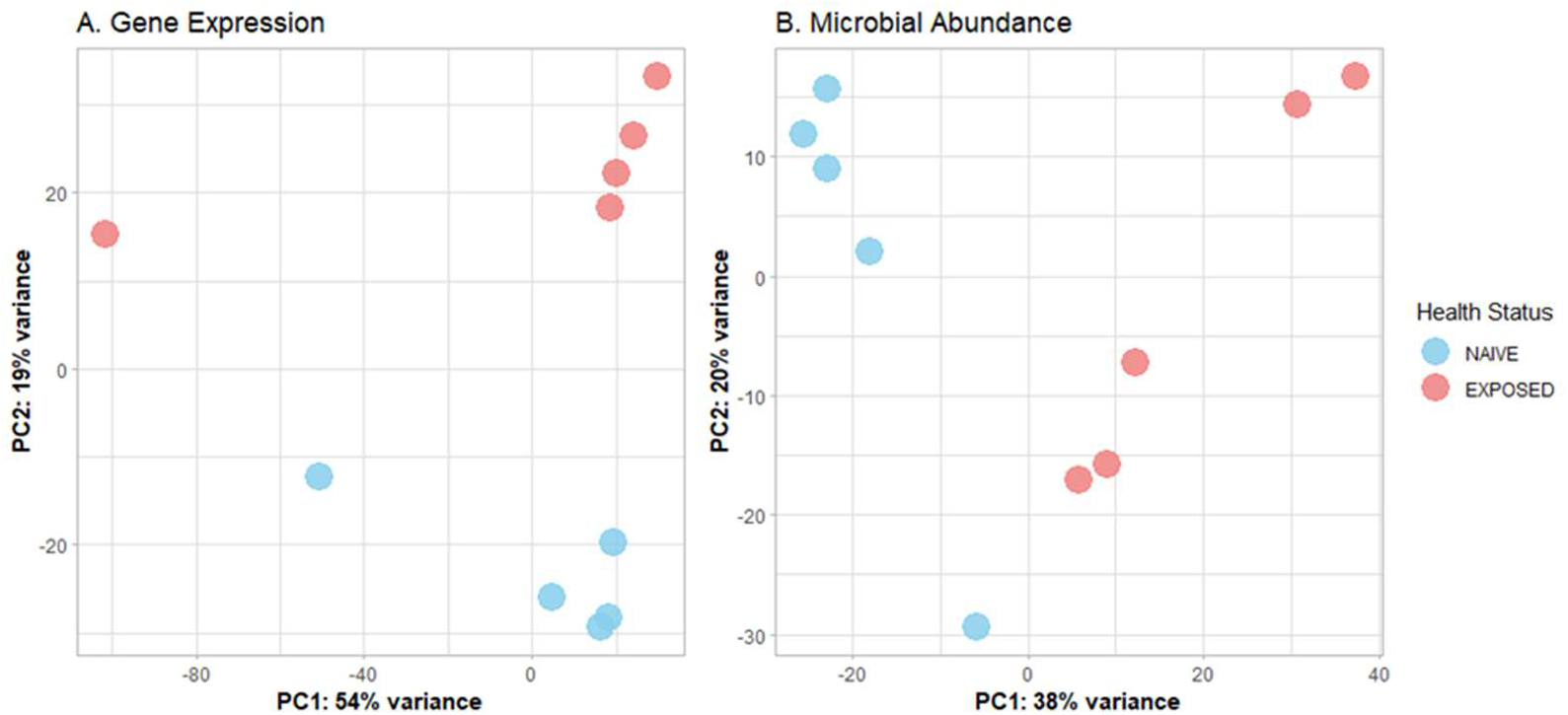
Principal component analysis of (**A**) gene expression, (**B**) microbial abundance. Blue circles represent naive individuals and orange dots represent exposed Individuals for gene expression and microbial abundance, respectively.

### Genomic Profiling revealed minimal Population Structure between sample sites

To test for population structure among our sampling locations, we ran PCA and admixture between naïve and exposed populations using the whole genome sequence data. We observed no significant distinction between sample sites in terms of population structure. Admixture was analyzed at multiple ancestry coefficients (*K*=1 to *K*=10). An ancestry coefficient of *K*=1 provided the best fit for the data with the lowest cross entropy score (***Figure S2.c)*.** In addition, at *K*=2, samples did not cluster by Naïve and Exposed sites (**Figure S2.b).** PCA was used to investigate genetic relationships in multivariate space. Naïve samples formed a distinct tight cluster while Exposed samples showed greater variation with no clear distinction between groups along either PC axis (**Figure *S2*.d**). Read Quality was verified between Naïve and Exposed groups with an average of 91.3% and 90.0% with Phread scores >30.

### Gene Ontology Enrichment (GO) reveals immune system, tissue integrity, stress response, and neurological changes in Exposed sea stars

To test for functional enrichment, we ran gene ontology analysis among genes differentially expressed between Naïve and Exposed sea stars. We found 249 enriched GO terms (*P* < 0.01) associated with health status, 130 of which were enriched among genes upregulated in the Exposed samples and 119 of which were enriched among genes upregulated in Naïve samples. After filtering out narrow and broad categories, and accounting for nested GO hierarchies, we found 73 and 48 unique terms enriched for upregulation in Exposed and Naïve groups, respectively (**Table S1-2**). A number of immune, cell stress, tissue integrity, and metabolism related categories were identified as enriched in the Exposed group and depleted in Naïve including humoral immune response (GO:0006959), amoeboid beta clearance (GO:0097242), response to toxic substances (GO:0009636), extracellular structure organization (GO:0043062), regulation of cell-cell adhesion (GO:0022407), protein folding (GO:0006457), response to hypoxia(GO:0001666), and various peptide and ATP metabolic processes (**Figure 2A, Table S1**). Fewer categories were enriched for upregulation in the Naïve group compared to Exposed, however a number of functions associated with nervous system function and cell stability were enriched including but not limited to dendrite development (GO:0016358), regulation of synapse structure (GO:0050803), cell adhesion (GO:0007155), cytoskeletal organization (GO:0007010), and neuromuscular junction development (GO:0007528) were enriched in the Naïve group (**Figure 2B, Table S2**) suggesting neural dysregulation in Exposed individuals. Using GO annotations to assign functional roles, we identified a total of 68 differentially expressed genes associated with immune system function, 79 in tissue development and homeostasis, 192 related to functions of the nervous system, and 144 involved in response to stress differentially expressed between Naïve and Exposed individuals (**Figure S3**).

**Figure 2.**
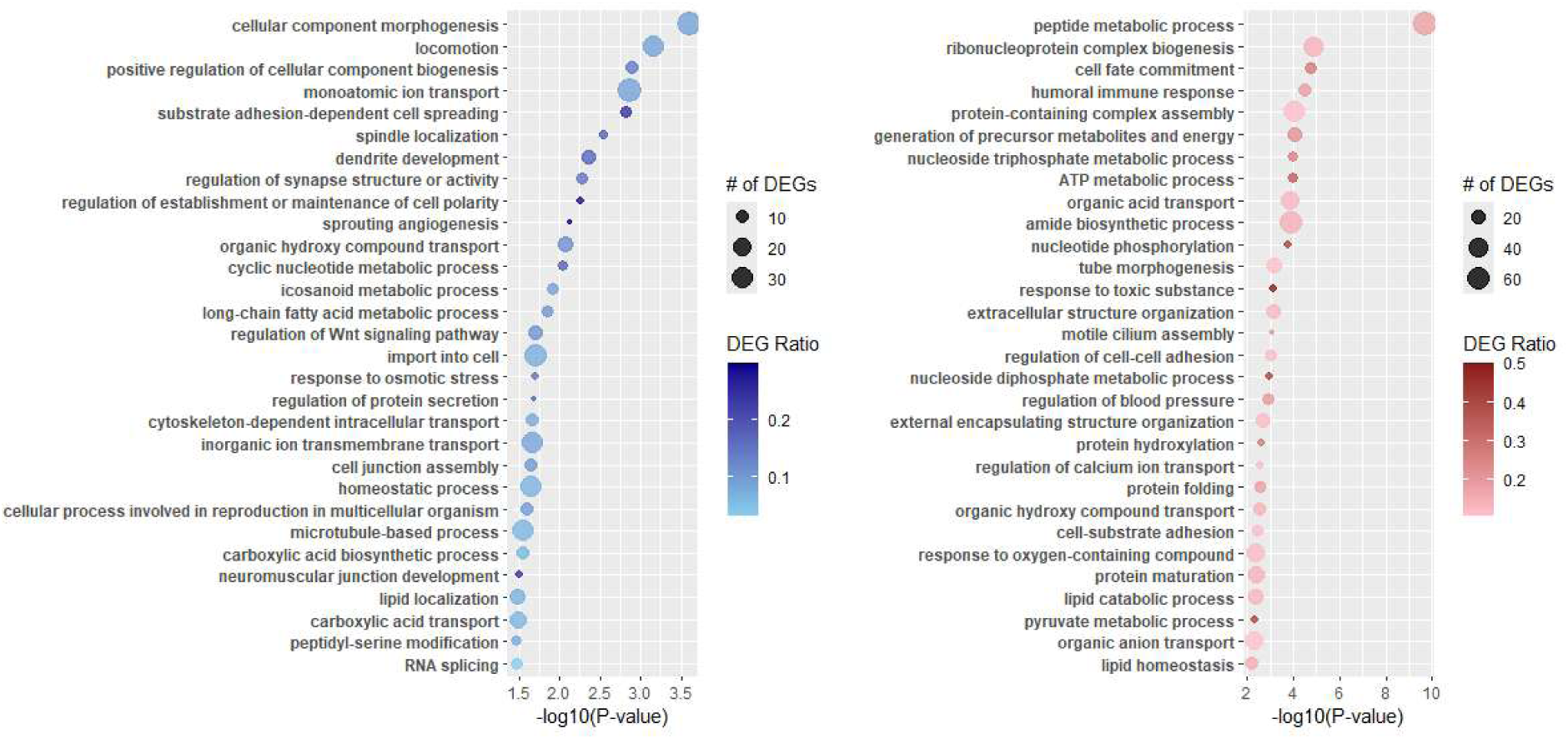
Gene Ontology enrichment plots for **A)** enriched for upregulation in Naive individuals compared to exposed and **B)** enriched for upregulation in Exposed compared to Naive. Top 30 enriched GO terms for naïve (blue) and exposed (red) health groups. X-axis represents the significance of enrichment transformed by -Log10 so that larger values represent more significant enrichment. All samples plotted have *P* < 0.01. Dot size represents the number of differentially expressed genes of the corresponding category that are differentially expressed. Color of each dot corresponds to the percentage of differentially expressed genes present in the category relative to all genes of said category.

### Functional annotation of differentially expressed genes

We compared gene transcripts against NCBI’s non-redundant protein database to assign functional annotations to differentially expressed genes. Of the 2006 differentially expressed genes, 1425 (78%) mapped to homologs, primarily in other sea star species; 381 (18%) mapped to uncharacterized homologs; and 80 (4%) had no annotation on the NCBI database. Our analysis revealed an activation of genes involved in the complement cascade, pathogen recognition, and variable expression in immune regulation and cell-death mediated pathways (**Figure 3A**). More specifically, we observed activation via upregulation of immune genes in Exposed stars including but not limited to *echinoidin*, *Ficolin-2-like*, *lymphocyte antigen 6-like* and *macrophage mannose receptor-1-like* (*MRC1*) and a decrease in tumor necrosis factors (TNF) family receptors. See Supplemental data on Dryad for a full list of differentially expressed genes and fold-changes (**DOI: 10.5061/dryad.t1g1jwtfh**:).

**Figure 3.**
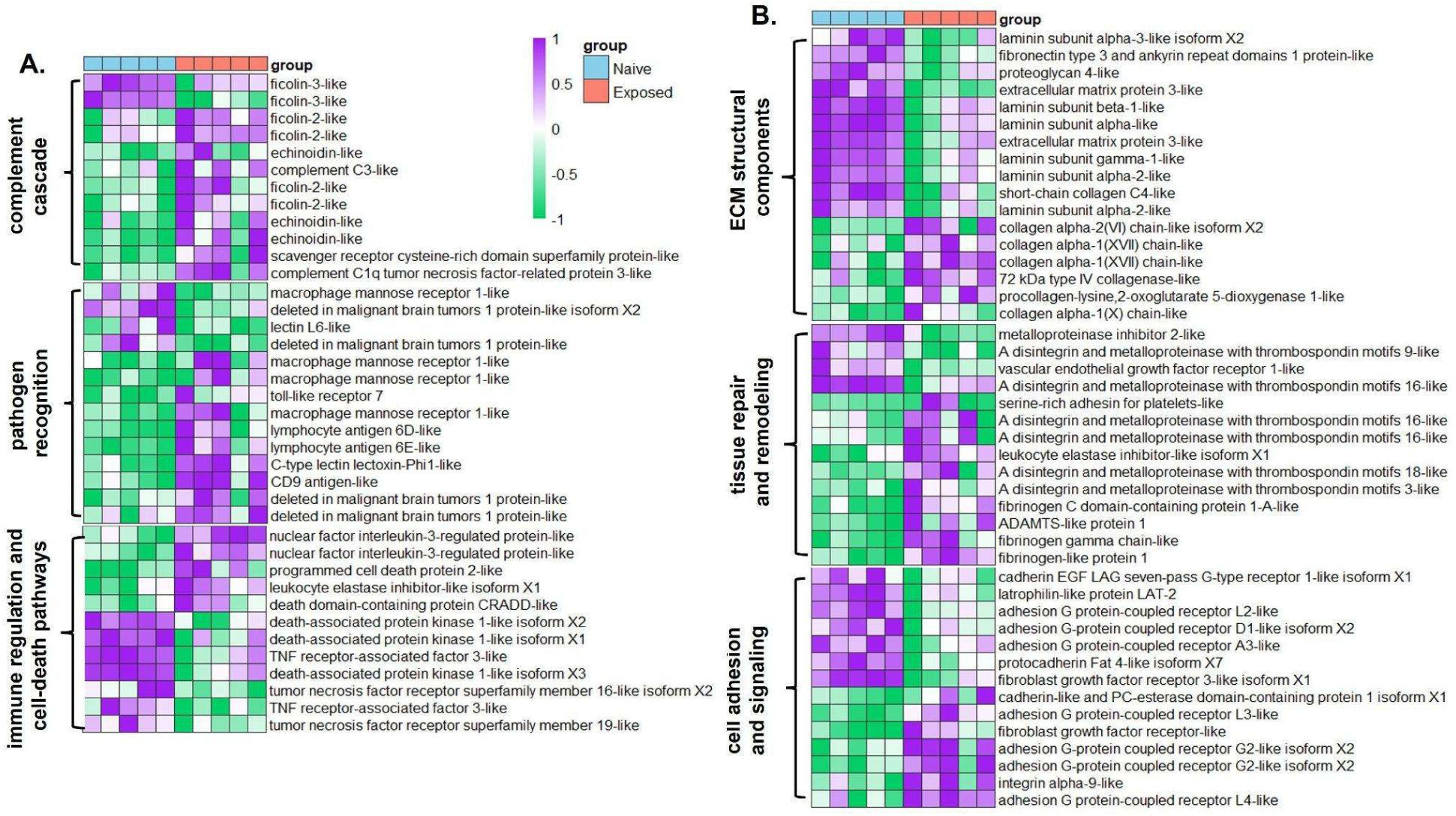
Differential Gene Expression Across Samples. Normalized gene expression patterns between naive (blue) and exposed (red) groups for differentially expressed genes with primary functions involved in **A)** immune system function and **B)** tissue integrity and remodeling. Higher expression values are shown in purple, lower expressions in green, and neutral in white.

Aligning with SSW symptoms (12, 22), we also document significant changes to tissue regeneration and homeostasis-related genes including extracellular matrix components, tissue repair and remodeling, and cell-adhesion and signaling genes (**Figure 3B**). The inflammatory immune response is closely tied to tissue integrity and homeostasis, highlighted here by variable expression in three fibrinogen genes that play a dual role as antimicrobial clotting agents and preservation of tissue integrity (56). Other genes involved in maintaining tissue integrity and essential for the breakdown and regeneration of extracellular structures were found with variable expressions including A Disintegrin and Metalloproteinase with Thrombospondin motifs (*ADAMTS*), *collagen alpha-like*, *short-chain collagen C4-like*, *Procollagen-lysine 2-oxoglutarate 5-dioxygenases-like (PLODs)*, *laminin* subunits-(𝛼*,β,γ*), and a number of adhesion G protein coupled receptors (aGPCRs) (**Figure 3B**).

Furthermore, we document a large number of genes associated with nervous system function and development showing differences in expression across health groups, some increasing, some decreasing in exposed versus naive samples, including multiple neuronal acetylcholine receptor subunit alpha genes, *neurotrypsin*, synaptotagmins, and *regulating synaptic membrane exocytosis protein 2* (*RIMS2*) (**Figure S4).** Finally, a number of cellular stress genes were expressed at higher levels in Exposed individuals including chaperon proteins *hsp60* and *hsp70*, *hsp90 co-chaperone Cdc37-like*, along with variable expression in detoxification genes NADPH-Oxidases and xanthine dehydrogenase/oxidase (**Figure S5).**

### Microbial Abundances differ between exposure groups

Three of the top five differentially abundant OTUs were of the family *Vibrionaceae*, including various unknown taxa mapping to the family *Vibrionaceae*, multiple species within the genus *Vibrio spp.,* and *Aliivibrio spp.* Other taxa of note that increased with exposure were the genus *Pseudoalteromonas*, *Colwellia* and *Photobacterium.* Among the top few OTUs less abundant among the Exposed group include the genus Chryseobacterium, *Sphingomonadaceae*, and *Thiobacillus* (**DOI: 10.5061/dryad.t1g1jwtfh**).

### Network analysis associates putative pathogenic microbes to immune response and tissue integrity

To utilize the power in variation among individuals, we used WGCNA to identify covarying gene expression and microbial abundance. Combined gene expression and microbial abundance data clustered into 39 distinct modules, 9 of which were significantly correlated with health status based on Pearson correlation between module eigengenes and trait values (*P* < 0.05; two-tailed *t*-test). To assess these relationships, we encoded health status as a binary numeric trait (i.e., Naïve = 0, Exposed = 1) and computed Pearson correlations between each module eigengene and this trait. Five modules showed a positive correlation—indicating higher eigengene values in the Exposed group—while four modules were negatively correlated, with higher values in the Naïve group. Of the 9 correlated with health status, 6 included both genes and microbes; of these, 4 had higher values for the Exposed group and 2 with higher values for Naïve (**Figure 4A**). These modules included 8,048 total genes and 45 microbial OTUs.

**Figure 4.**
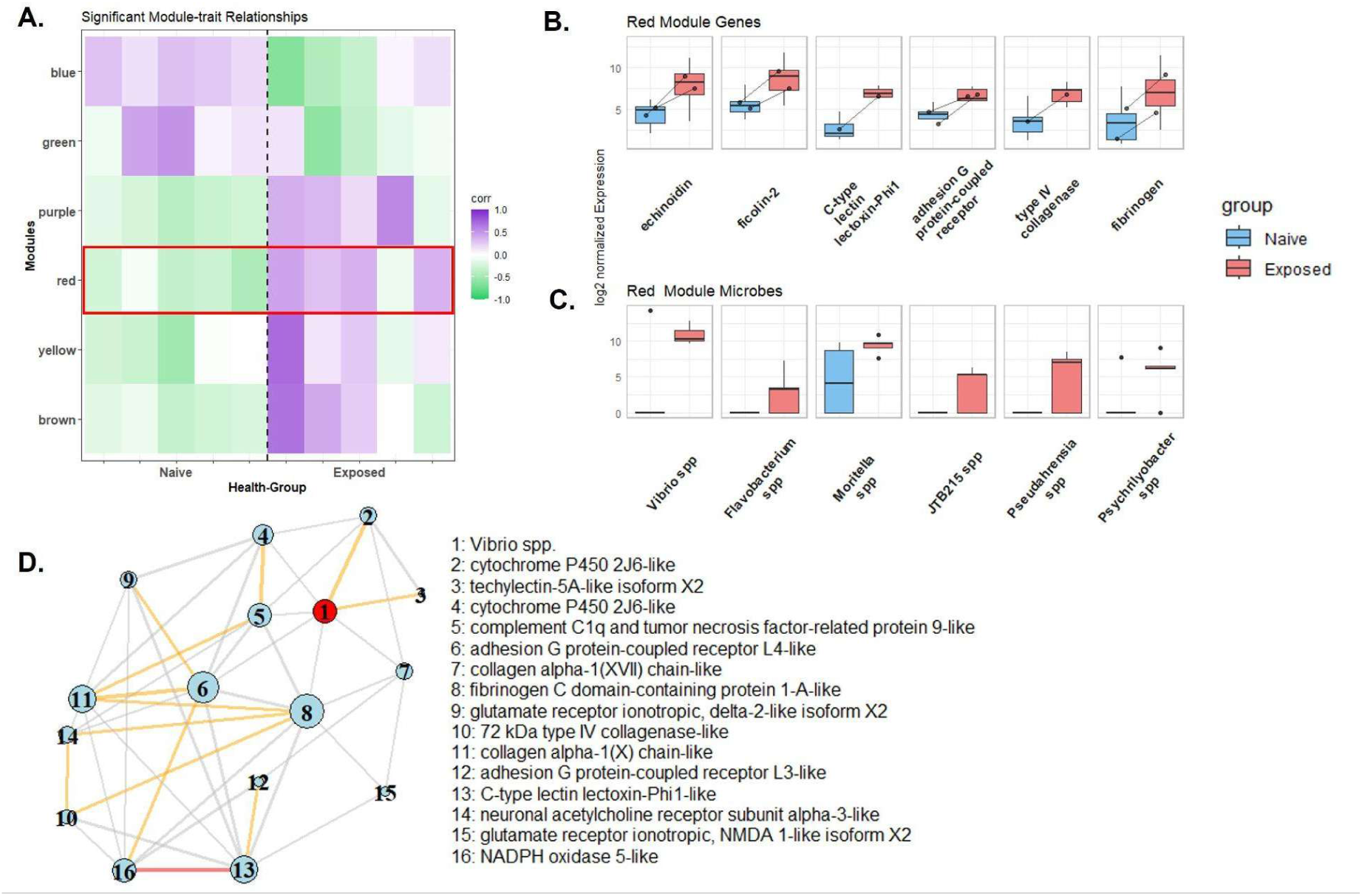
**A**). WGCNA correlation matrix across samples for modules containing both genes and microbes with positive correlation (purple) and negative correlation (green) **B**) log-2 expression plots between naive (blue) and exposed (pink) groups of select genes in the “Red” module. **C**) log-2 abundance plots between naive (blue) and exposed (orange-red) groups of select microbes in the “Red” module. **D)** Network plot of neighboring genes (sharing an edge) to *Vibrio spp.* with correlation value > 0.5 of genes involved in immune, tissue homeostasis, detoxification, and neural related genes. Red edges represent correlation values above 0.9, orange above 0.7, and grey above 0.5.

The “Red” module (see red box in **Figure 4A**) showed the strongest association with the treatment group (*P* = 0.001; two-tailed *t*-test), with the highest number of differentially abundant microbes (9) and genes (770) in any significant module containing both genes and microbes. The “Red” module contained some of the most differentially abundant microbes in our analysis, including OTUs mapping to *Vibrio spp., Flavobacterium spp., Moritella spp., JTB215 spp., and Pseudahrensia spp (***Figure 4C)**. Of the 770 genes in the “Red” module, 346 (45%) were differentially expressed including many within categories of interest relative to signs associated with SSW including immune function and host defense (fibrinogens, lectins, echinodin, ficolins, *lymphocyte antigen 6*, *c-type lectin*), collagen degradation and synthesis (*collagen alpha-1(XVII) chain-like*, type IV collagenase-like, ADAMTS, aGPCRs), DNA repair, and nervous system function (neuronal acetylcholine receptor subunits; **Figure 4B).** Gene ontology analysis of the red module revealed biological functions related to immune system process, cell-cell adhesion, response to wounding, and response to oxygen levels (**Figure S6A**).

Due to our previous observations of a dramatic increase in abundance of *Vibrio* at the first onset and just prior to visible signs of wasting (27, 29) and the strong correlations with *Vibrio* in the present transcriptomics data, we further explored the “Red” network by pruning nodes (genes and microbes) and edges (correlation values) to focus on the strongest connections with *Vibrio* species. We filtered to keep only genes with enriched annotations, *i.e.*, those associated with immune response, tissue integrity, neural function, and stress response, and keeping edges with correlations > |0.5|. *Vibrio spp.,* showed direct correlations with host expression of techylectin 5A (involved in pathogen recognition), *fibrinogen C*, collagen alpha subunits, *adhesion G protein coupled receptor L4*, and *complement C1q facto*, and *cytochrome p450* genes with secondary connections to C-type lectin lectoxin-Phi1, and type IV collagenase (**Figure 4D**).

The “Purple” module was the second most significant between health group status (*P* = 0.001; two-tailed t-test) containing 246 total genes, and eight OTUs including *Pseudoaltermonas spp., Vibrionaceae spp., and Aliivibrio spp.* which were among the differentially abundant taxa. Of the 246 genes, 124 were among those differentially abundant, including multiple heat shock proteins (*hsp70*, *hsp90*), *fibrinogen-like protein 1*, *procollagen-lysine*, and xanthene dehydrogenase/oxidase proteins. Gene ontology analysis of the purple module revealed enrichment in categories mainly involved in stress response including protein folding and metabolic processes (**Figure S6B**).

## Discussion

In this study, we analyzed samples from wild *Pycnopodia helianthoides* undergoing their first recorded outbreak of SSW in this region. Most SSW research has focused on symptomatic individuals within controlled lab settings, limiting insights into disease progression in natural environments. By comparing asymptomatic sea stars from yet-unimpacted and active SSW areas, we capture a critical and rarely studied early stage of disease. Through integrated transcriptomic and microbiome analyses, we detected immune, nervous, and collagen system changes strongly correlated with increased abundances of putative pathogenic microbes isolated from epidermal tissue. This result has important implications, as multiple studies have documented a rapid accumulation of opportunistic bacteria once symptoms are visible, potentially obscuring responses specifically to the initial causative agent(s). By capturing this critical time point, our analysis helps illuminate core immune mechanisms in response to SSW exposure in the wild, refining our understanding of the disease’s progression.

We highlight two important findings. First, we observed immune system activation of sea stars exposed to SSW in their direct environment with relatively higher expression of immune genes compared to individuals sampled from sites not affected by the outbreak. This observation is consistent with the hypothesis that nervous system and tissue-integrity dysregulation precede the loss of neuromuscular and collagenous tissue control that are commonly observed in wasting sea stars (12). This hypothesis is further supported by the reduced and variable expression of collagen, cell adhesion, cell-signaling, neural development and synaptic-component genes in exposed individuals. We find further evidence of cellular stress through the activation of genes coding for DNA repair, heat-shock proteins, and oxidative stress pathways.

Second, we find evidence of host-microbe interactions dependent on disease exposure. This insight was revealed by leveraging network analyses integrating host gene expression data with microbiome data from the same tissue biopsies. Notably, changes in host transcript levels and microbial abundances were correlated across samples in relation to health status. Indeed, microbes that were identified in these modules were previously associated with exposure and health status and are consistent with previous studies investigating changes to microbiome composition associated with SSW (27–30). Furthermore, the genes connected through these same modules are primarily involved in immune response, maintaining tissue integrity, cell adhesion, metabolism, and nervous system processes. Our detection of immune activation, stress, and dysregulation of neural and tissue homeostasis in conjunction with shifts in putatively pathogenic microbial taxa in asymptomatic (yet exposed) individuals builds upon existing evidence of immune activation during SSW and further implicates the role of microbial interactions as a key factor, if not driver, of disease progression.

The activation of immune mechanisms has been documented in laboratory exposures to SSW in both *P. helianthoides* and *Pisaster ochraceus* (26, 30, 38). However, these previous studies investigated previously Exposed individuals compared to those actively showing signs of wasting. Here, we find evidence of similar pathway activation between unexposed and exposed individuals to SSW as previous studies found between exposed and actively wasting individuals. Fuess *et al.* identified several immune, neural, and tissue-regulation pathways in lab-Exposed *P. helianthoides* documenting differential expression in humoral immune factors—particularly cytokines, lectins, and components of the complement system (ficolins, macrophage mannose receptors, and lectins) align with an expected immune mobilization against pathogenic threats. Specifically, we found strong increases in lectins such as *echinoidin*, an echinoderm specific lectin that plays a role in the defense system against microorganisms (57), and *lectin L6-like* which aggregates and limits growth around Gram-positive and Gram-negative bacteria (58, 59) accompanied by a strong increase in *ficolin-2-like* genes responsible for self-non-self-recognition (60). The strong degree of increase in immune genes observed in wild populations provides new insights, as it reveals immune responses initiated in the field before visible symptoms appear. Interestingly, unlike Fuess *et al*., we did not detect significant changes in toll mediated pathways expression in our samples, potentially due to the epidermal nature of our tissues sampled in comparison to their direct sampling of immune cells (coelomocytes) which would likely carry a higher level of pattern recognition receptors (61).

Differential regulation in neural and cell-adhesion genes was not unexpected. A large number of neuronal and tissue related genes can respond to disease or injury as animal nervous and immune systems cooperate to assess and respond to danger (62). Neurons play a vital role in defense response, including regulating immune function, healing injuries, and pathogen recognition (63) and can be mediate cell adhesion and connective tissue, including collagenous tissue essential for maintaining sea star structures (64). In our analysis, we discovered evidence of such interactions. Among numerous cell-signaling related genes, seven total *adhesion G-Protein Coupled Receptors* (aGPCRs) were found to be differentially regulated. aGPCRs are known for their roles in regulating cell-cell adhesion and transmitting signals between the extracellular matrix and intracellular components (65). aGPCRs ability to interact and bind with ECM components highlights concurrent changes found in extracellular matrix remodeling genes including a number of collagen alpha chain subunits, *type IV collagenase*, *short-chain collagen 4*, and *A disintegrin and metalloproteinase (*ADAMTS) have been shown to be essential for tissue regeneration in the sea cucumber *Holothuria glaberrima* (66, 67). Changes to these signaling and structural components in the extracellular matrix may contribute to the “melting” phenotype often associated with sea stars suffering from SSW alongside the loss of control of their catch-collagen system. Proper regulation of collagen homeostasis may help animals resist wasting in some species of sea stars, supported by increased collagen gene expression in *P. ochraceous* that resist wasting compared to those that succumb to disease (30). This is sensible as more rugose (soft) species, such as *P. helianthoides* which has one of the highest mortality rates on record, show lower levels of SSW resistance than more *“hard and smooth”* species such as the bat star, *Patiria miniata* (28, 68).

Network analysis revealed clustering of specific bacterial taxa, notably *Vibrio spp.*, with genes linked to immune, tissue integrity, and stress responses, implicating facultative anaerobes in early disease interactions. Multiple studies have detected the proliferation of *Vibrio*, *Pseudoalteromonas*, and other facultative anaerobes in association with SSW disease progression (26–30). Gudenkauf and Hewson 2015 were the first to investigate SSW through a metatranscriptomic lens, discovering increase in *Pseudomonas* and *Vibrio* relatives coinciding with apoptotic, tissue degradation and signaling of death-related processes (26). Lloyd & Pespeni 2018 highlighted a progressive change in microbiome composition though disease stages of *P. ochraceous* including *Vibrio* and *Pseudoalteromonas,* and later Aquino et al., 2020 implicated the role of facultative anaerobes at the animal-water interface accumulating with disease progression (27, 28). The original microbial data used in this analysis was collected from a subset of those analyzed in McCracken et al., 2023, which further characterized changes to the microbiome before visible signs were present among more individuals of wild *P. helianthoides* (47 Naïve, 20 Exposed, 18 Wasting). In the previous study, we found an increase of *Vibrio* species by ∼1200 fold, along with other facultative anaerobes upon disease exposure (29). While this signal was reduced to ∼7-fold increase in the subset of samples for the present study (5 Naïve and 5 Exposed), the genus *Vibrio* retained the highest change of abundance. Furthermore, we found direct connections between *Vibrio spp.* and a C-type lectin lectoxin, where C-type lectin expression has been previously associated with anti-vibrio infection in copepods (69).

It is well established that *Vibrio spp.* can cause disease in both vertebrates and invertebrates, including vibriosis in fish, cholera in humans caused by *Vibrio cholerae*, and lethal infections through open wounds from exposure to pathogenic *Vibrio* in marine environments (70, 71). Members of the genus *Vibrio* are of particular interest to this study as they have been implicated in a number of historic disease outbreaks impacting other marine species including echinoderms. One study investigating population control strategies in the crown of thorns star, *Acanthaster planci,* were able to induce strikingly similar symptoms to SSW by injecting agar that promotes *Vibrio* growth which was then transmissible among and between species of sea stars (33, 34, 72). Many *Vibrios* have been implicated in other echinoderm diseases including bald urchin disease and black spot disease (31, 73) with strikingly similar SSW-like symptomology characterized by fast-progression skin lesions and tissue necrosis. *Vibro’s* are known to cause disease in a host of other marine species, including *V. coralliilyticus* which is known to cause tissue damage and necrosis in corals (74), and *Vibrio parahaemolyticus* involved in Skin Ulceration Disease (SKUD) impacting the sea cucumber *Holothuria scabra* (32).

The causes of bacterial infections can be difficult to identify and often include interacting biotic and abiotic factors to align. The parallels across studies linking the proliferation of facultative anaerobes increased immune activity, and transcripts suggestive of low-oxygen stress in both *P. ochraceous* and *P. helianthoides* support a model where rising sea temperatures and localized hypoxia, while not required for disease, may foster ideal environmental conditions for SSW to initiate and spread. In support of this hypothesis, Pespeni and Lloyd 2023 discovered a higher expression of hypoxia related genes (*HIF-1*) in individuals that seemed to resist wasting as compared to those that succumbed to wasting (30). Although we did not detect changes specifically in *HIF-1*, our analysis revealed the enrichment of genes involved in response to oxygen containing compounds, and response to hypoxia (**Table S2**). Furthermore, thermal stress such as those caused by rising sea surface temperatures, has been shown to have direct impacts on immune system function in purple sea urchins, *Strongylocentrotus purpuratus,* decreasing the antibacterial efficiency of their immune cells (75). We detected increased expression of a number of heat-shock proteins in our analysis, however they also may be responding as part of a general cellular-stress pathway associated with infection. Other marine bacterial diseases, such as Pacific Oyster Mortality syndrome (PCOM) on oyster populations, and SKUD can be triggered by environmental factors via the injection or ingestion of organic matter and changes in environmental temperature (32, 76). These complicated biotic-abiotic interactions may explain why SSW was not a major threat to sea star communities until relatively recent years as sea temperatures continue to rise.

## Data availability statement

The datasets presented in this study can be found in online repositories. The names of the repository/repositories and accession number(s) can be found below: BioProject, *PRJNA1286911*. Dryad: DOI: 10.5061/dryad.t1g1jwtfh.

## Author contributions

All authors contributed to and approved the manuscript. Conception and design: AM, MP. Data Generation: AM. Data Analysis: AM,AR,CS,SS,SB,MP. Visualizations: AM. Writing & Editing: AM, JCBN, MP.

---

**Figure S1:**
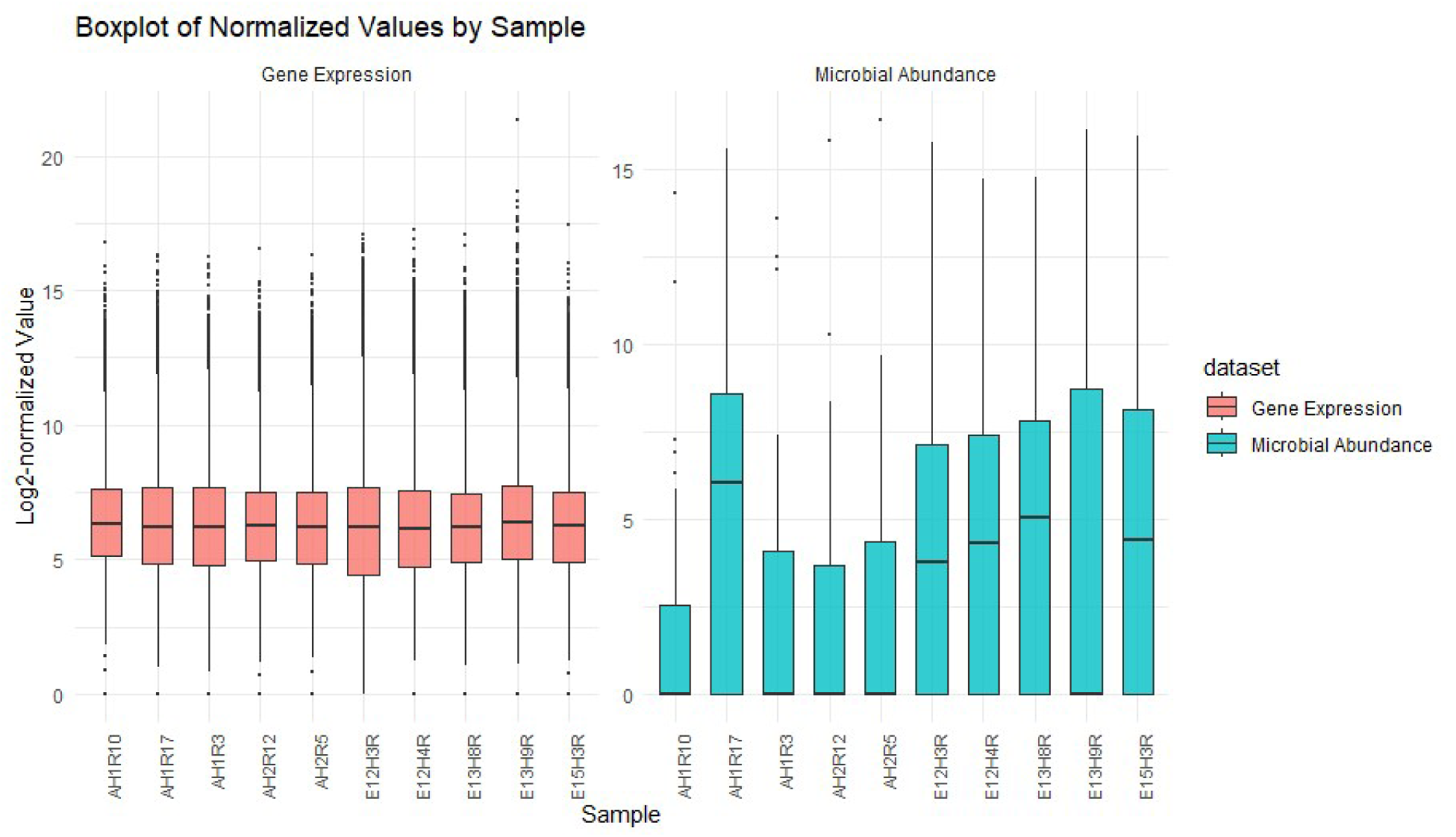
Log-2 normalized count distributions per sample for gene expression (red-pink), and microbial abundance (turquoise) used in joint WGCNA analysis.

**Figure S2:**
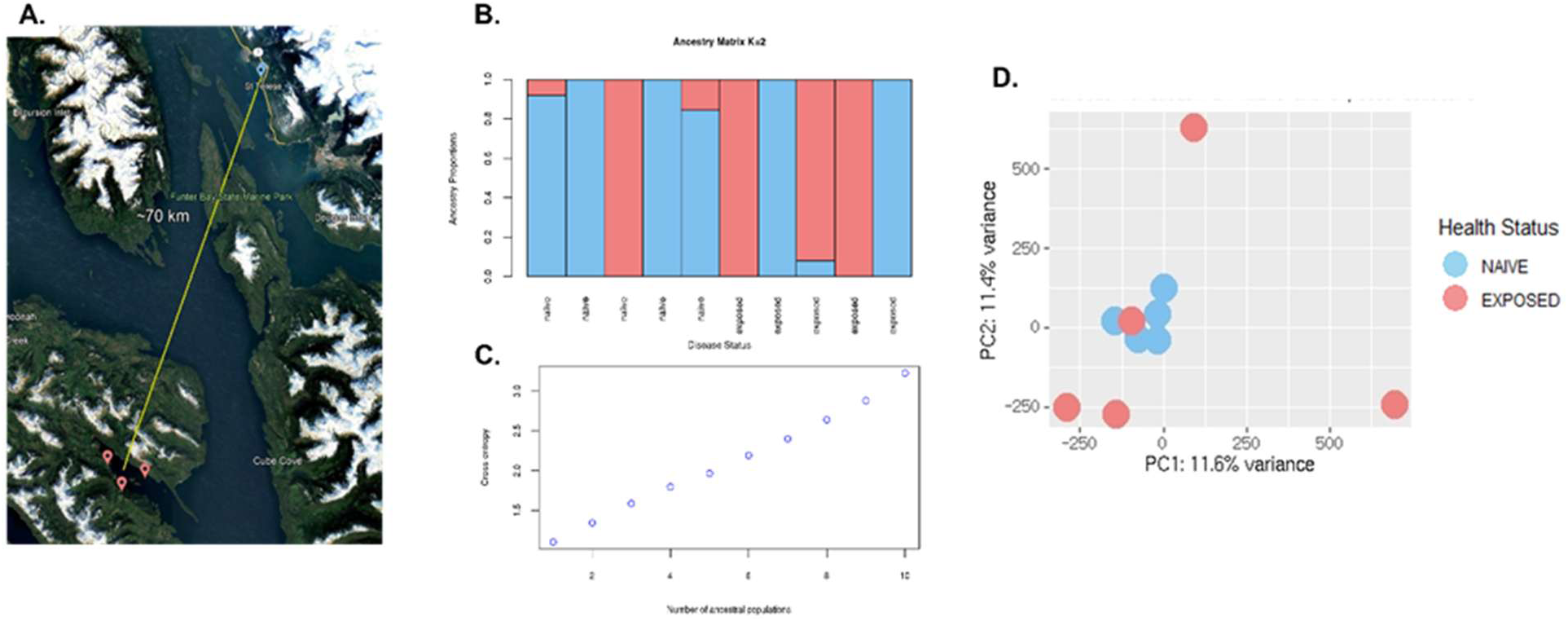
Population genetics analysis of site-health groups. **A)** locations of sample sites. **B)** Admixture plot K=2, **C)** Ancestry Coefficients, **D)** principal component analysis of single nucleotide polymorphisms between Naive (blue) and Exposed (pink) individuals.

**Figure S3:**
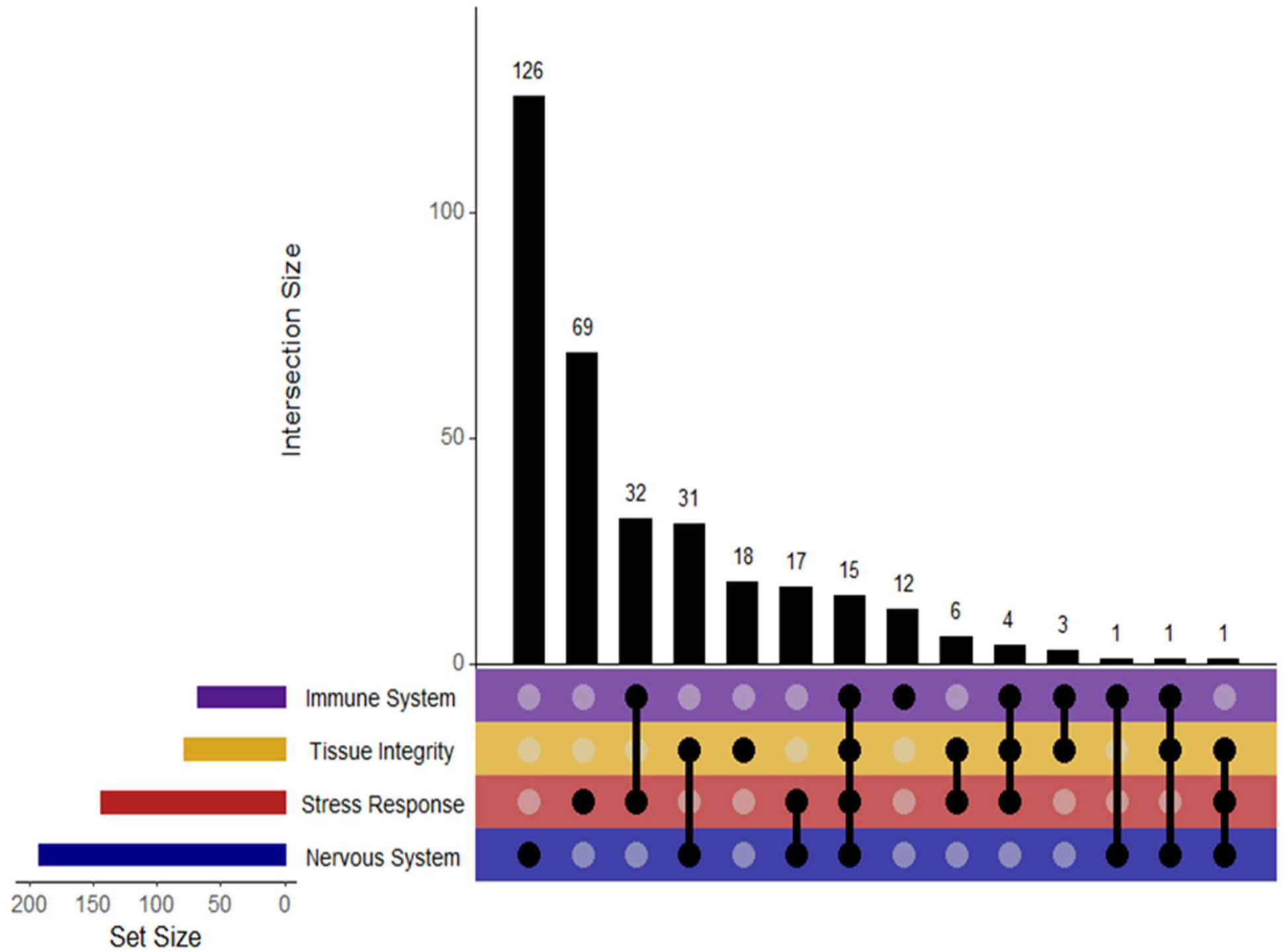
Number of differentially expressed genes mapping to Gene ontology categories with the following functions: Immune System Function (Purple), Tissue Integrity and Homeostasis (Yellow), Cellular and organismal stress response (Red). Nervous System Function (Blue). Hight of the bars indicate the number of genes in each category where filled in circles in each colored row indicate functional category.

**Figure S4.**
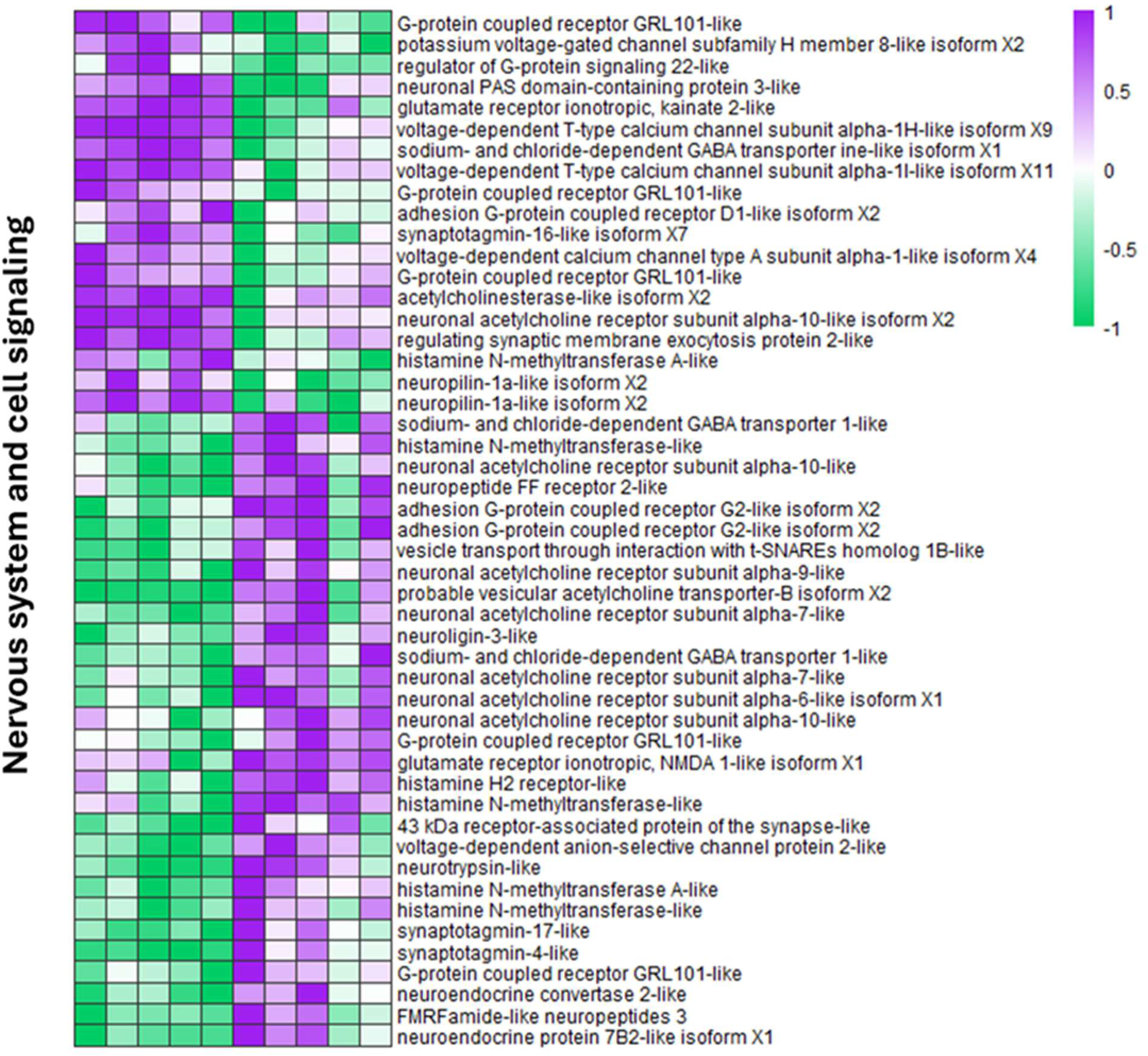
Differential Gene Expression Across Nervous System genes. Normalized gene expression patterns between Naive (blue) and Exposed (red) groups for differentially expressed genes with primary functions involved in Nervous System Function. Higher expression values are shown in purple, lower expressions in green, and neutral in white.

**Figure S5.**
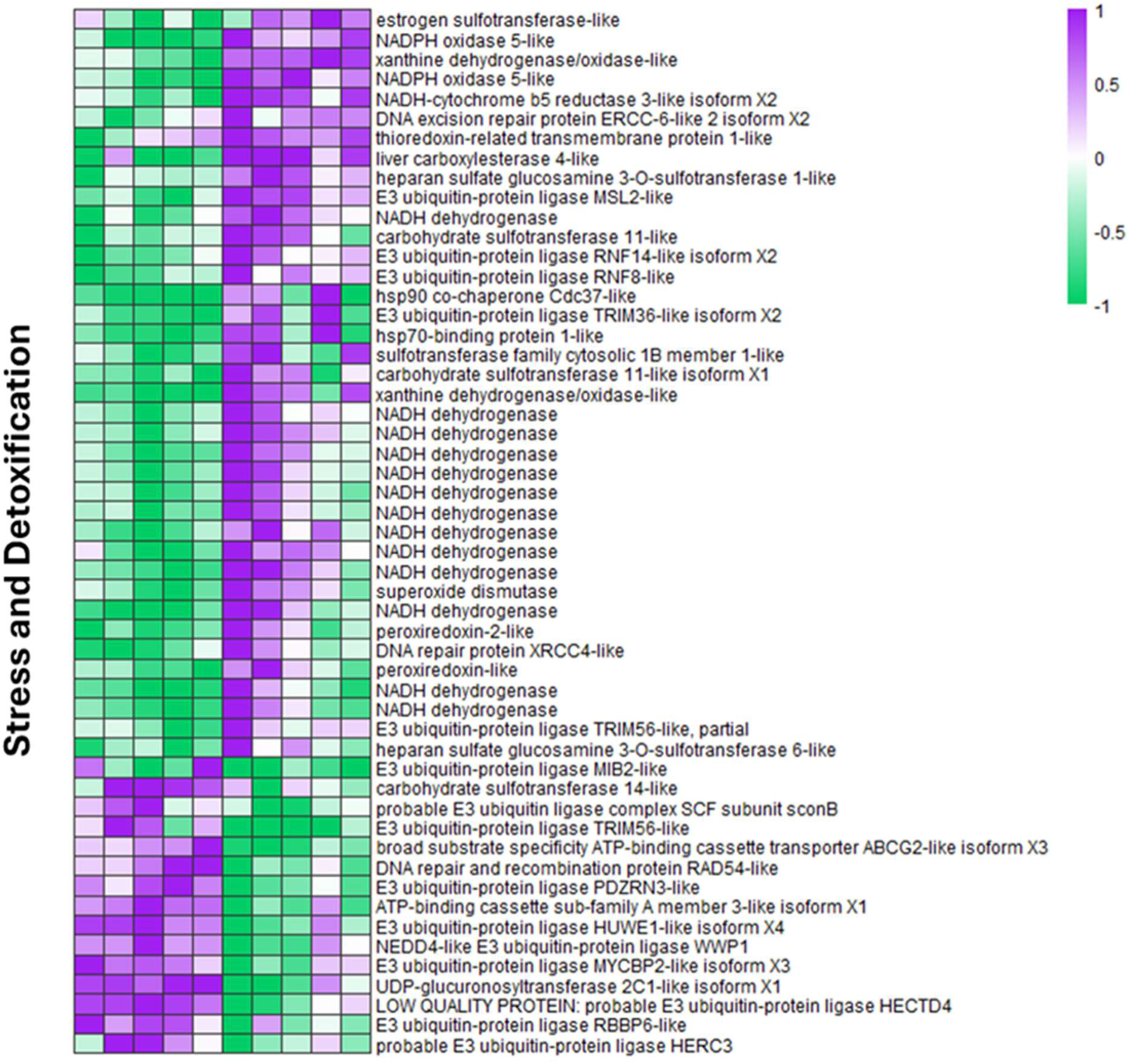
Differential Gene Expression Across Stress and Detoxification Genes. Normalized gene expression patterns between Naive (blue) and Exposed (red) groups for differentially expressed genes with primary functions involved in stress and detoxification. Higher expression values are shown in purple, lower expressions in green, and neutral in white.

**Figure S6:**
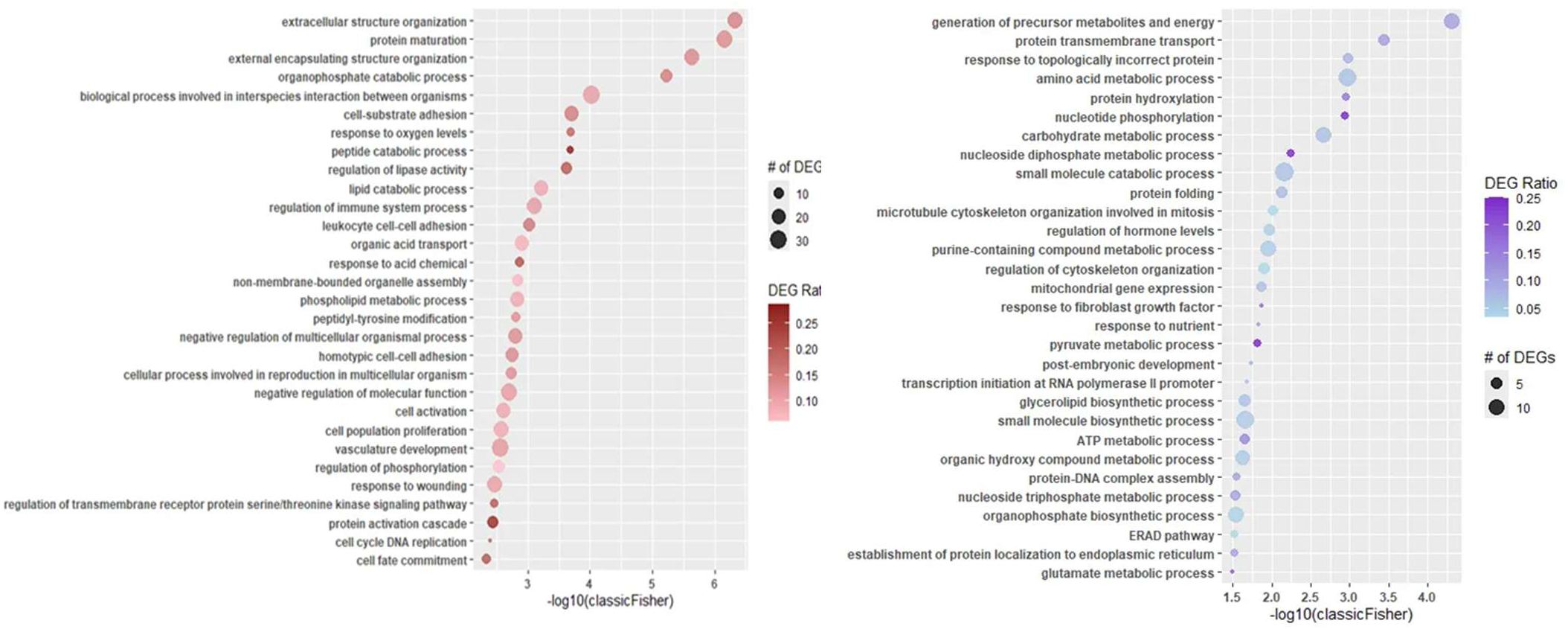
WGCNA Module Gene Ontology Enrichment Plots. A) Enriched for upregulation in Red Moduleand **B)** enriched for upregulation in Purple Module. X-axis represents the significance of enrichment transformed by -log10 so that larger values represent more significant enrichment. All samples plotted have p<0.01. Dot size represents the number of differentially clustered genes appearing in each GO category from the WGCNA module. Color of each dot corresponds to the percentage of differentially expressed genes present in the category relative to all genes of said category

